# Aromatic lactic acid production by *Bifidobacterium longum* subsp. *infantis* is determined by cell density, substrate availability and pH *in vitro*

**DOI:** 10.64898/2025.12.17.694872

**Authors:** Thilde Garbøl Plenge, Mikael Pedersen, Martin Frederik Laursen

## Abstract

*Bifidobacterium longum* subsp. *infantis (B. infantis)* is an important colonizer of the infant gut. This species is known to produce a variety of health beneficial metabolites, including aromatic lactic acids (ALAs), supporting early life immune development. However, the regulation of ALA production in *B. infantis* remains poorly understood. In this study, we investigated how environmental factors, such as substrate availability and pH affect *B. infantis* DSM20088 growth and ALA production *in vitro*. Bacterial batch cultivations supplemented with different amino acid concentrations revealed a linear relationship between indole-3-lactic acid (ILA), 4-hydroxyphenyllactic acid (OH-PLA), and phenyllactic acid (PLA) production and exogenous concentrations of tryptophan, tyrosine and phenylalanine, respectively. Furthermore, chemostat cultivations at physiological relevant pH levels (4.5, 5.5 and 6.5) showed that an acidic pH slowed growth and shifted ALA production by *B. infantis*. Overall, ALA concentrations correlated strongly with bacterial abundance across and within pH levels, showing that cell density is a major determinant of ALA production. After considering the growth limiting effects of lower environmental pH, it had limited impact on the total production of ALAs, but the relative production of PLA was stepwise enhanced at more acidic pH at the expense of ILA, independent of cell density. In conclusion, this study shows that cell density, substrate availability and environmental pH determine aromatic lactic acid production by *B. infantis*.

**Importance:** *Bifidobacterium longum* subsp. *infantis* is an abundant bacterial species in the infant gut microbiota where it supports immune development e.g. though the production of aromatic lactic acids. In this study we cultured *Bifidobacterium longum* subsp. *infantis* at different concentrations of aromatic amino acids and pH levels to investigate the effect of these environmental factors on aromatic lactic acid production. We show that there is a linear relationship between aromatic lactic acid production and the exogenous aromatic amino acid concentration, that the production strongly follows cell density, and that pH regulates the preference towards producing different aromatic lactic acids. This knowledge may help inform strategies to enhance or direct beneficial ALA production in the infant gut.

## Introduction

‘Infant-type’ *Bifidobacterium* species are important members of the early life gut microbiota. These species are promoted by breastfeeding due to their ability to metabolize human milk oligosaccharides (HMOs), the third-most abundant solid component in breastmilk [1]. *Bifidobacterium longum* subsp. *infantis* (*B. infantis*) in particular, has a prodigious capacity to grow on HMOs [2] and dominate the gut ecosystem in breastfed infants [3]. Moreover, high abundance of *B. infantis* contributes to beneficial immune maturation, reduced intestinal inflammation, and improved intestinal barrier integrity [4].

One way by which *B. infantis* contributes to host physiology is through the production of metabolites, including aromatic lactic acids (ALAs) [5]. ALAs are derived from the aromatic amino acids (AAAs), tryptophan, tyrosine and phenylalanine, which are metabolized to indole-3-lactic acid (ILA), phenyllactic acid (PLA), and 4-hydroxyphenyllactic acid (OH-PLA), respectively. It has been shown that ALAs aid in immune regulation by serving as ligands for host cell receptors [6], [7]. Especially, ILA is recognized for its ability to activate the aryl hydrocarbon receptor (AhR), which is involved in epithelial barrier function, protection against pathogens and regulation of host metabolism [8], [9]. Additionally, ILA and PLA are potent ligands for the hydroxycarboxylic acid receptor 3 (HCA_3_), a regulator of immune functions and energy homeostasis [5], [10], [11]. ALAs also serve as antimicrobial agents [12], [13], [14] and has been suggested to reduce the prevalence of *E. coli* and other antimicrobial resistance gene-rich taxa in the infant gut [15].

*B. infantis* produces ALAs via a catabolic pathway that likely begins with the transamination of AAAs into aromatic pyruvic acids, which are subsequently converted to ALAs by the aromatic lactate dehydrogenase Aldh [4]. While it has been shown that ‘infant-type’ *Bifidobacterium* species, including *B. infantis* produce ALA in the infant gut [5], how the gut environment affects the accumulation of these metabolites remains unknown.

Physical and chemical factors in the intestinal environment such as nutrient availability, pH and transit time all influence the composition and metabolic activity of the microbiota [16], [17]. Infants are usually fed breastmilk, formula, or a combination of both, each providing different substrates available for bacterial fermentation in the gut. Although commercial formula is designed to mimic human milk, most formulas contain no or limited amounts/types of HMOs, leading to a lower abundance of ‘infant-type’ *Bifidobacterium* species [18]. Additionally, the AAA content in human milk changes during the lactation period and differ between formula brands [19]. Since AAA concentrations direct ALA production by lactic acid bacteria [20], amino acid availability might play a role in ALA production by *B. infantis*. Moreover, correlations between the abundance of ‘infant-type’ *Bifidobacterium* species and ALA concentrations in infant faeces have been observed [5], suggesting that higher cell densities of producer-species result in higher production of ALAs. However, since high abundance of *Bifidobacterium* species, in particular *B. infantis*, is also negatively correlated to faecal pH [21], a lower gastrointestinal pH, favouring ALA production, might also explain the higher concentrations of ALAs. Supporting this, it has been reported that acid stress leads to upregulation of genes involved in amino acid metabolism in *B. longum* subsp. *longum* [22], highlighting the possibility that pH regulates ALA production. We therefore investigated how AAA substrate availability, cell density and environmental pH affect ALA production by *B. infantis* DSM20088.

## Results

### ALA production follows AAA availability in a dose dependent manner

To test the effect of amino acid availability on ALA production, batch cultures of *B. infantis* were grown in MRS-cys broth supplemented with different concentrations of AAAs. After 48 h of growth, cell density and metabolite accumulation were assessed. All cultures reached an optical density (OD_600nm_) between 2.6 - 3.6 (**Supplementary Figure 1.A**). The cultures with the highest amino acid supplementation (0.2%) grew to a slightly, but significantly lower OD_600nm_ than non-supplemented cultures, indicating that the starting concentration of AAAs or accumulating metabolites slightly limit final cell densities under these conditions.

ALAs and AAA concentrations were assessed by LC-MS. ILA, OH-PLA and PLA were not detected in the MRS-cys broth control. In inoculated cultures, explicit production of these metabolites was observed (**Figure 1.A**). PLA was the most abundant ALA in the non-supplemented culture (208±29 μM), followed by OH-PLA (105±13 μM) and ILA (50±8 μM). This is in line with phenylalanine being major aromatic amino acid in the MRS-cys broth (**Supplementary Figure 1.C**) and have been reported previously [5].

**Figure 1:**
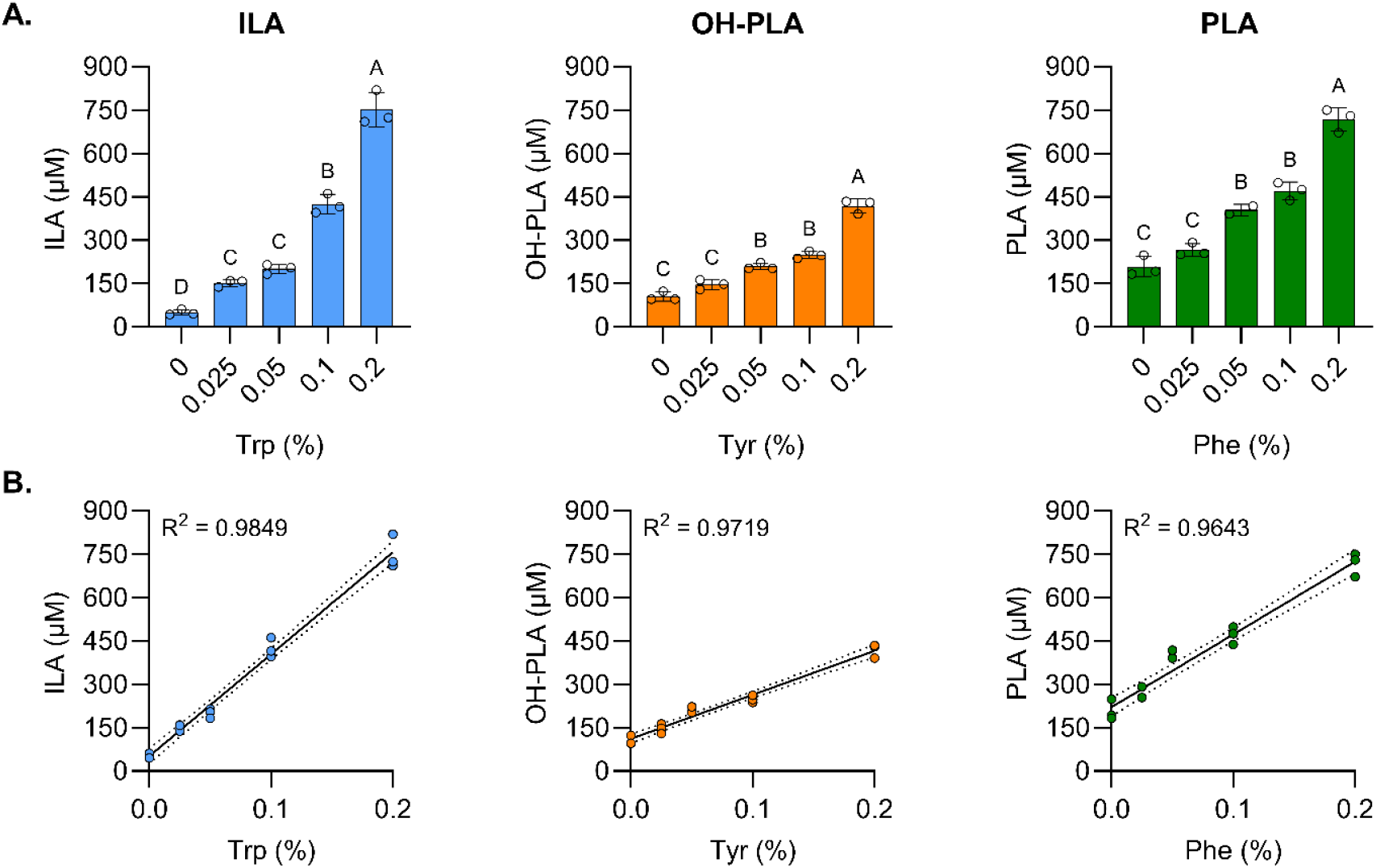
Aromatic lactic acid concentrations (μM) in B. infantis batch cultures grown for 48h in MRS-cys medium with aromatic amino acid supplementation. **(A)** Increase in aromatic lactic acid concentration in response to supplementation. Bars show mean ± s.d. of three biological replicates. Statistical significance was evaluated by one-way ANOVA followed by Tukey’s multiple comparisons test. Means not sharing letters are significantly different with adjusted p < 0.05. **(B)** Proportional relationship between aromatic lactic acid concentration and percent supplementation with R_2_-values shown in the figure. Points (n = 15) show a single biological replicate. Abbreviations: ILA = indole-3-lactic acid; PLA = Phenyllactic acid; OH-PLA = 4-hydroxyphenyllactic acid; Trp = tryptophan; Phe = phenylalanine; Tyr = tyrosine.

In tryptophan, tyrosine and phenylalanine supplemented cultures, a significant increase in ILA, OH-PLA and PLA was observed, respectively (**Figure 1.A**). Moreover, the production of each ALA showed a strong linear relationship with the percent of supplemented AAA (**Figure 1.B**). ILA and PLA showed clear proportional relationships and very strong positive correlations with the detected tryptophan and phenylalanine concentrations, respectively (**Supplementary Figure 1.B**). OH-PLA did not correlate with measured tyrosine concentrations, but this result is suspected to be caused by technical issues arising from the lower solubility of tyrosine.

### ALA production is tightly coupled to cell density

To investigate the effect of different physiological pH levels on ALA production, *B. infantis* was cultivated in chemostat cultures maintained at constant pH levels of 4.5, 5.5 and 6.5 (**Figure 2.A**). The cultures were grown for 24 hours in batch mode and thereafter in chemostat mode until 96 hours (**Figure 2.B**). For pH 6.5 and 5.5 cultures, there was no significant difference in OD_600nm_ over time (**Supplementary Figure 2.A**). Additionally, these cultures grew with almost the same growth rate (**Figure 2.C**). pH 5.5 cultures maintained a constant OD_600nm_ from 24 hours and onward, while pH 6.5 cultures OD_600nm_ stabilized after 48 hours (**Figure 2.B**). Growth was to some extent inhibited at pH 4.5 since both the growth rate (**Figure 2.C**) and OD_600nm_ over time (**Supplementary Figure 2.A**) were significantly lower than for the less acidic pH conditions. pH 4.5 cultures did not show a stabilization in OD_600nm_ when applying chemostat mode (**Figure 2.B**). For the first 24 hours of chemostat mode, the pH 4.5 culture continued to increase and peaked at the 48-hour timepoint. Hereafter a decrease in OD_600nm_ was observed. Consistently, CFU counting over time showed a significant decrease in viable cell numbers only at pH 4.5 between 48-96 h, whereas cell numbers were stable kept at ≈2.4x10^9 CFU mL−1 for pH 5.5 and ≈1.3x10^9 CFU mL−1 for 6.5 between 48-96h (**Supplementary Figure 2.B**). As expected, CFU counts over time were significantly correlated with OD_600nm_, supporting the growth patterns of *B. infantis* observed in OD_600nm_ measurements across the different pH conditions (**Supplementary Figure 2.C**).

**Figure 2:**
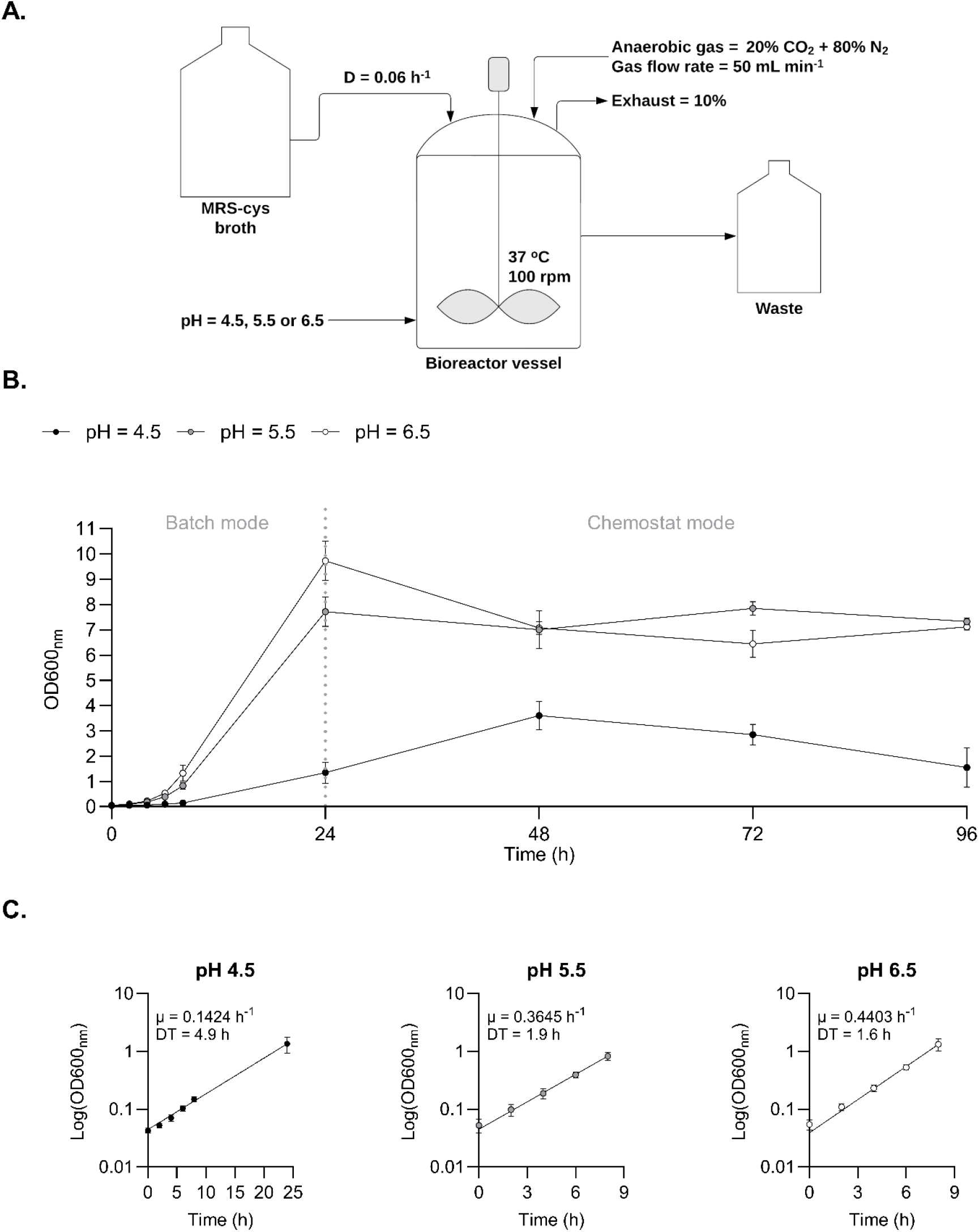
Chemostat cultures of B. infantis at different pH levels. **(A)** Chemostat culture parameters. **(B)** Optical densities (OD_600nm_) over time. The cultures were run in batch mode for the first 24 h whereafter chemostat mode was applied. Points show mean ± s.d. of three biological replicates. **(C)** Log transformed optical densities (OD_600nm_) of B. infantis during the exponential growth phase with the specific growth rate (μ) and doubling time (DT) at each pH level. Points show mean ± s.d. of three biological replicates.

Total and individual ALA concentrations largely followed OD_600nm_ measurements over time within each pH condition (**Figure 3.A**). In general, higher ALA concentrations were observed for pH 5.5 and 6.5 compared to pH 4.5 (**Supplementary Figure 3**), mirroring the growth differences. Across pH conditions, ALA concentrations correlated positively with OD_600nm_ (**Figure 3.B**) and CFU mL^-1^ of *B. infantis* (**Supplementary Figure 4**). At the individual pH level, pH 4.5 and 6.5 also showed significant positive correlations between OD_600nm_ and ALAs (**Supplementary Figure 5**). This was not observed in pH 5.5 cultures, possibly explained by steady state in cell density already after 24 h. Overall, this shows that the production of ALAs is tightly coupled to the cell density of *B. infantis*.

**Figure 3:**
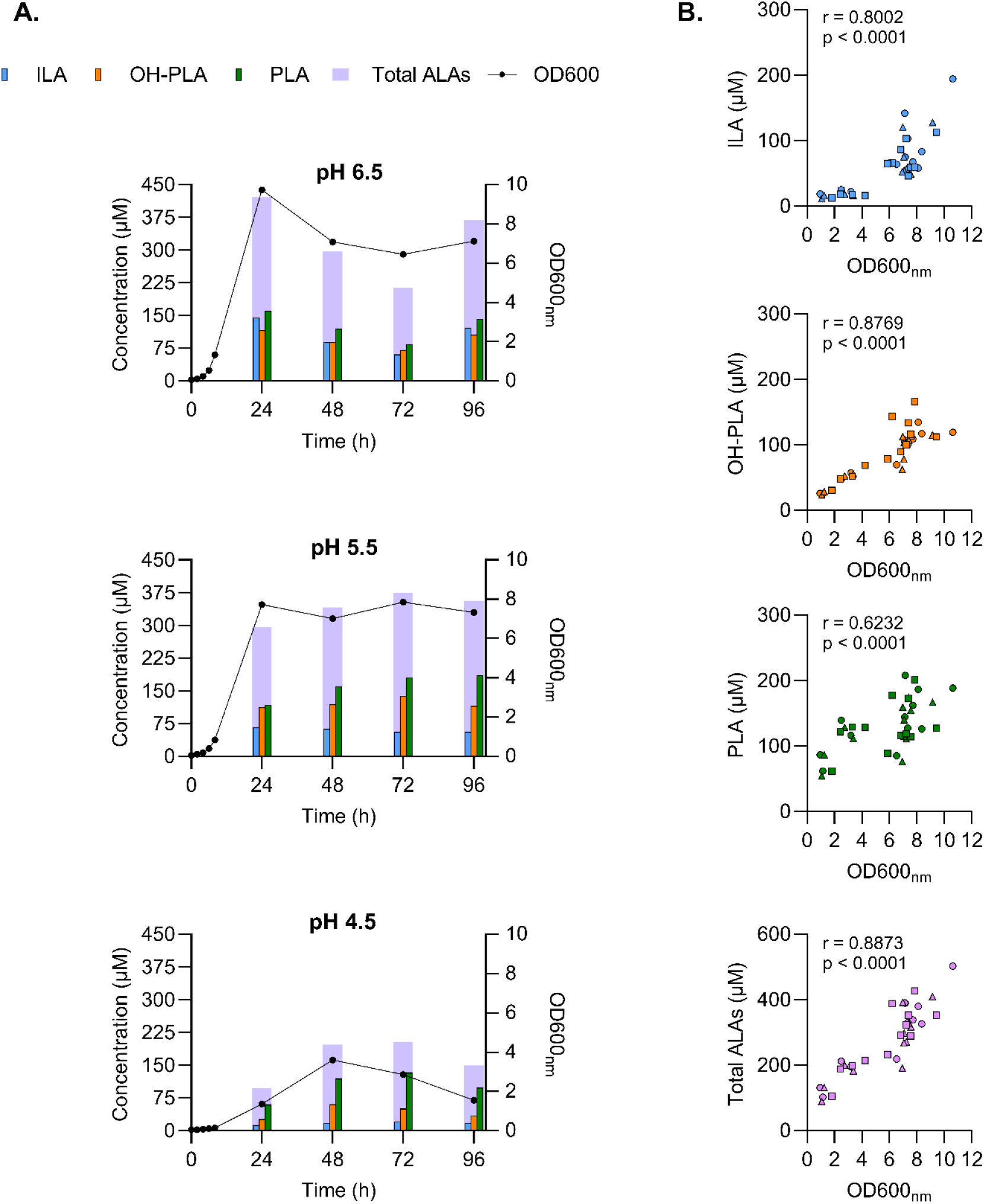
Aromatic lactic acid production and growth of B. infantis in chemostat cultures at different pH levels. **(A)** ILA, OH-PLA, PLA, total ALAs concentrations (μM) and optical densities (OD_600nm_) over time. Bars and points show means of three biological replicates. **(B)** Correlations (n = 36) between optical density (OD_600nm_) and ALAs (μM) in chemostat cultures at 24-96 h with Pearson correlation coefficients (r) and p-values (two-tailed) shown in the figures. Biological replicates across pH levels are shown as ▴, ▮ and ?. Abbreviations: ILA = indole-3-lactic acid; PLA = Phenyllactic acid; OH-PLA = 4-hydroxyphenyllactic acid; ALAs = aromatic lactic acids.

### pH has limited influence on total ALA production but regulates preference towards the type of ALA produced

While ILA, OH-PLA and total ALA concentrations were lower at pH 4.5, PLA production did not differ between pH 4.5 and 6.5 (**Supplementary Figure 3**). However, PLA levels were lower at both pH 4.5 and 6.5 relative to pH 5.5. Additionally, OH-PLA concentrations were elevated at pH 5.5, while ILA levels were significantly higher at pH 6.5. No significant difference in total ALA concentrations was observed between pH 5.5 and 6.5.

Since cell density had a pronounced impact on ALA production and varied across different pH levels, ALA concentrations were normalized to OD_600nm_. Normalized concentrations showed a significant, but minor influence of pH on ALA production (**Figure 4**). ILA exhibited the highest normalized production at pH 6.5, though this increase was only statistically significant relative to pH 5.5. The normalized OH-PLA levels were elevated at pH 4.5 compared to pH 6.5. Additionally, normalized PLA concentrations were higher at both pH 4.5 and pH 5.5 compared to pH 6.5. The total normalized production of ALAs was significantly higher at pH 4.5 than at the other tested pH conditions, mostly driven by the elevated PLA production. However, since there were no differences in normalized total ALA production between pH 5.5 and 6.5, and limited difference to pH 4.5 (somewhat driven by the decrease in cell density over time under this condition), it appears that pH regulates the preference towards producing different ALAs rather than the total production. To investigate this further, the relative percentwise abundance of the three ALA at the different timepoints were compared (**Figure 5**). At pH 6.5, an almost equal abundance of the three ALAs was observed. Oppositely, when lowering the pH, a stepwise shift towards higher PLA and less ILA production occurred. Therefore, taking into account its effect on growth, environmental pH has limited influence on the total production of ALAs but rather seem to regulate the preference towards the type of ALA produced.

**Figure 4:**
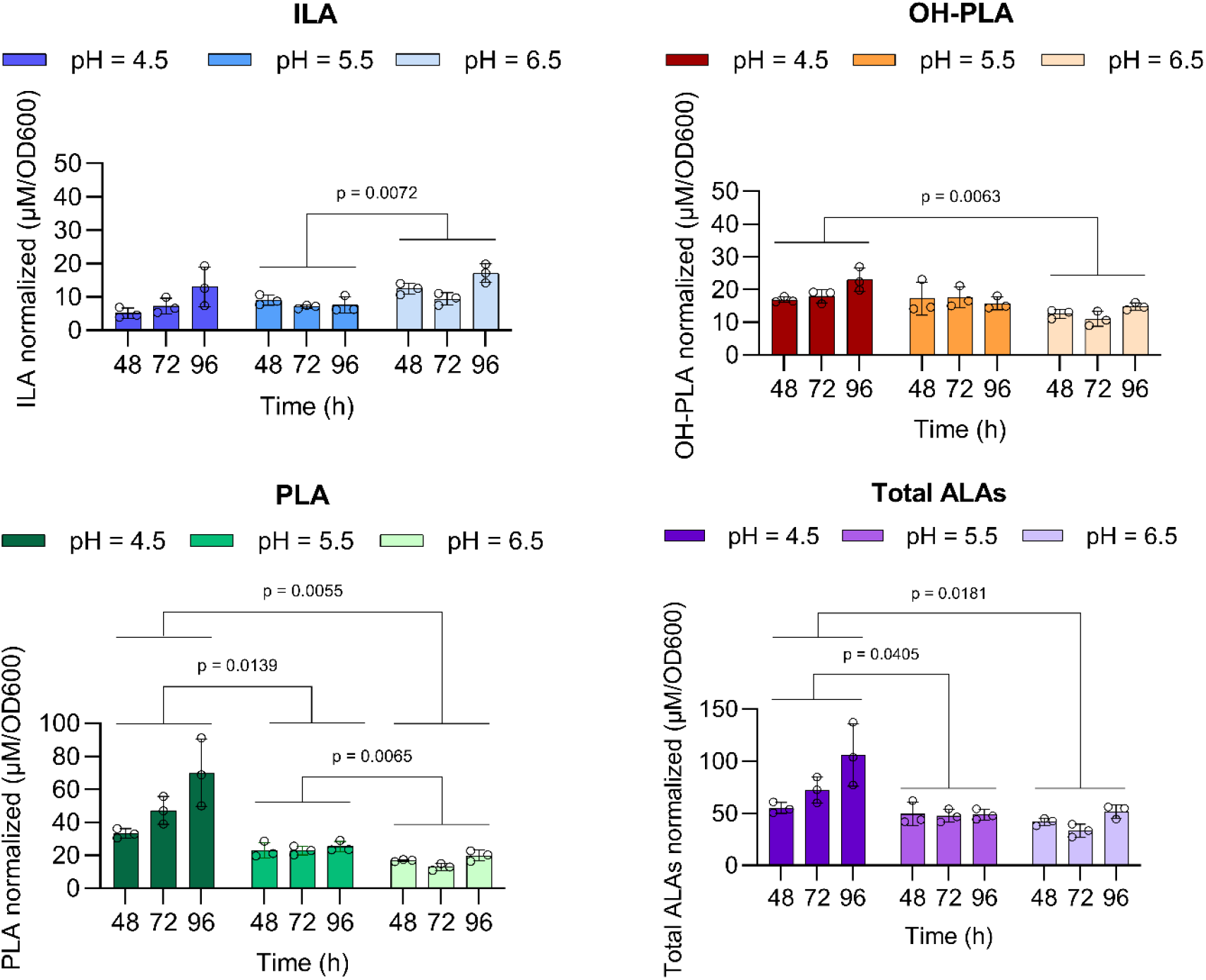
Normalized aromatic lactic acid production by B. infantis grown in chemostat cultures at different pH levels during steady state. Aromatic lactic acid concentrations normalized with OD_600nm_ (μM/OD600) at different pH levels. Bars show mean ± s.d. of three biological replicates. Statistical significance between groups were evaluated with two-way ANOVA with p-values of the pH effect shown on the figures. Abbreviations: ILA = indole-3-lactic acid; PLA = Phenyllactic acid; OH-PLA = 4-hydroxyphenyllactic acid; ALAs = aromatic lactic acids.

**Figure 5:**
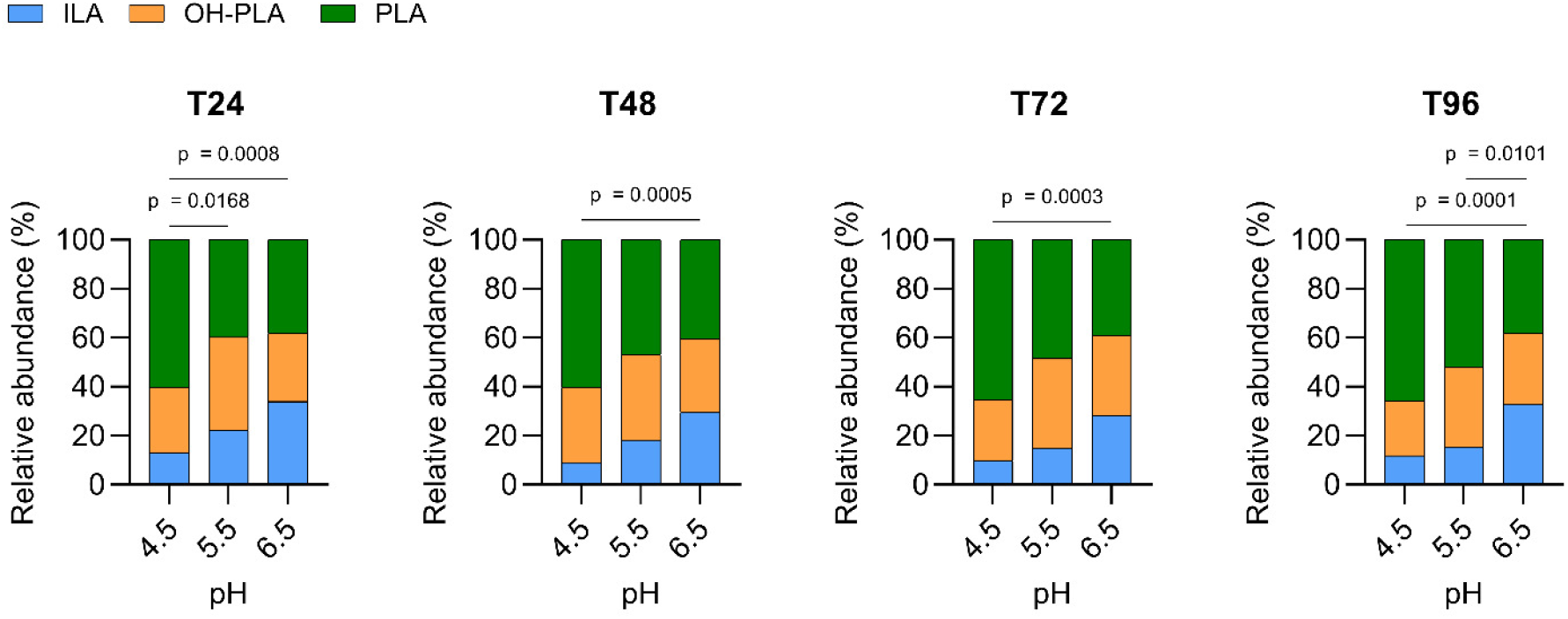
Preference in aromatic lactic acid production by B. infantis grown in chemostat cultures at different pH levels. Relative distributions (%) of aromatic lactic acids at different timepoints. Stacked bars show the means of three biological replicates. Statistical significance was assessed using the Chi-squared test with p-values displayed in the figure panels. Abbreviations: ILA = indole-3-lactic acid; PLA = Phenyllactic acid; OH-PLA = 4-hydroxyphenyllactic acid; ALAs = aromatic lactic acids.

### Acidic pH conditions lead to subtle upregulation of *aldh* expression and affect several *aat* genes

The *aldh* gene is responsible for the last step in the formation of ALAs by ‘infant’-type *Bifidobacterium* species [5]. To get a deeper understanding of pH regulated ALA production, we therefore quantified *aldh* (BLON_RS05510) expression by reverse transcription quantitative polymerase chain reaction (RT-qPCR). It was indicated that an acidic pH leads to a slight upregulation of *aldh* (**Figure 6.A**), consistent with the slightly higher normalized ALA concentrations at pH 4.5 (**Figure 4**). However, since the major effect of pH was a shift in the type of ALA being produced, we hypothesized that other genes involved in aromatic amino acid metabolism might exhibit more dramatic pH-driven changes. Therefore, RNA sequencing (RNAseq) was conducted to assess global transcription. In agreement with the RT-qPCR, RNAseq also showed a slight upregulation of the *aldh* gene, especially at pH 4.5 (**Figure 6.B**).

**Figure 6:**
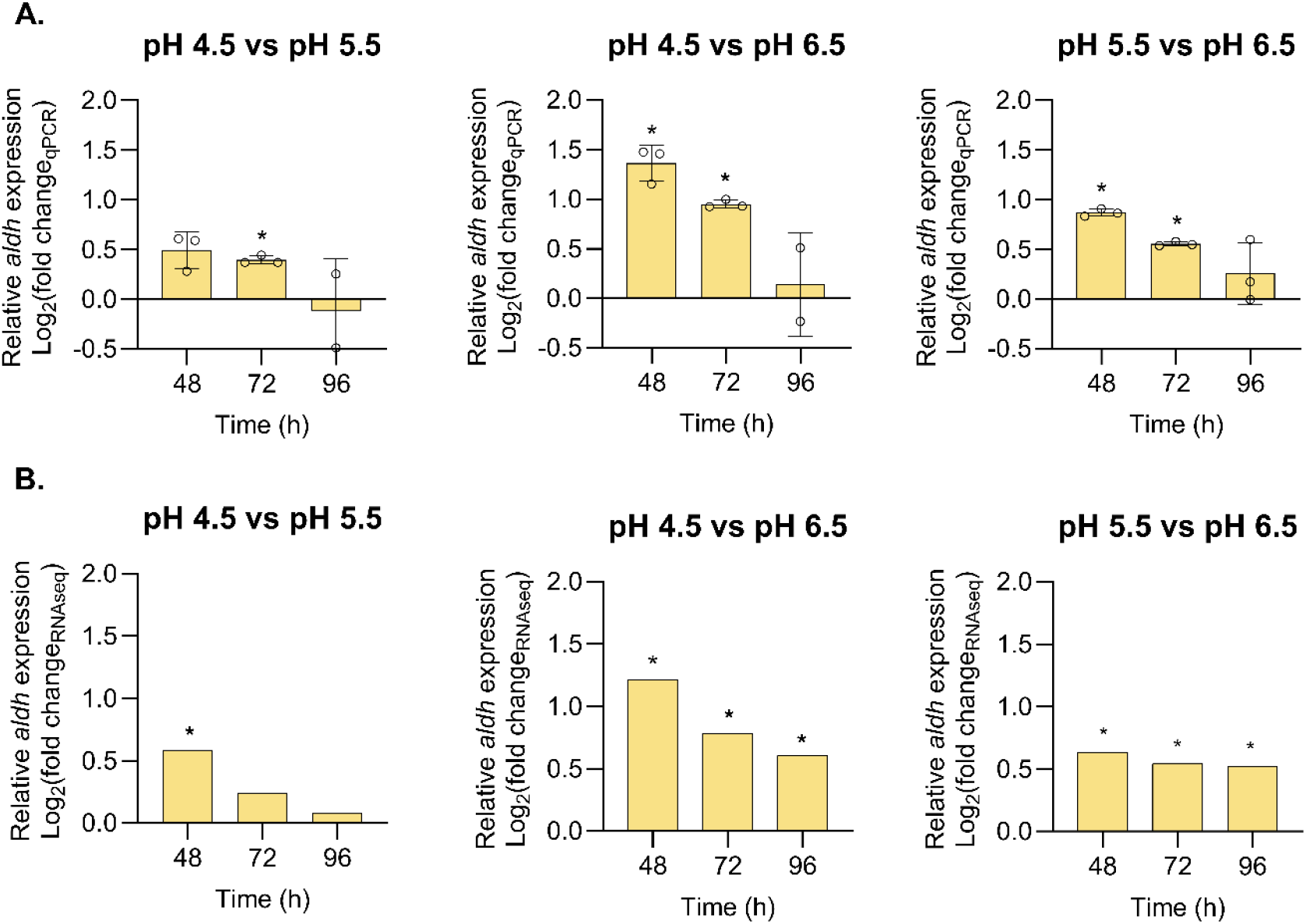
Relative aldh (BLON_RS05510) expression between pH conditions. **A)** Log_2_ fold change of the relative expression determined by RT-qPCR. Bars show mean ± s.d. of three biological replicates. Statistical significance was assessed using an unpaired two-tailed t-test with Welch’s correction on gene expression values normalized to reference genes. * p < 0.05. **B)** Log2 fold changes at each time point, calculated from three biological replicates. *: Genes with an average read count greater than 100 across all samples and an FDR-corrected p-value < 0.1 for the log2 fold change are considered significant.

Because *aldh* is located within a genetic element that contains a haloacid dehalogenase gene (*had*) (BLON_RS05515) and an amino-acid transaminase gene (*aat*) (BLON_RS05520), we examined the expression patterns of these neighboring genes as well (**Table 1**). The *had* gene followed the expression pattern of *aldh* and showed a small upregulation at pH 4.5 and 5.5 compared to pH 6.5. Interestingly, the *aat* gene was on the other hand consistently more than 2-fold upregulated at pH 4.5 compared to the less acidic conditions with no differential expression observed between pH 5.5 and 6.5.

**Table 1:**
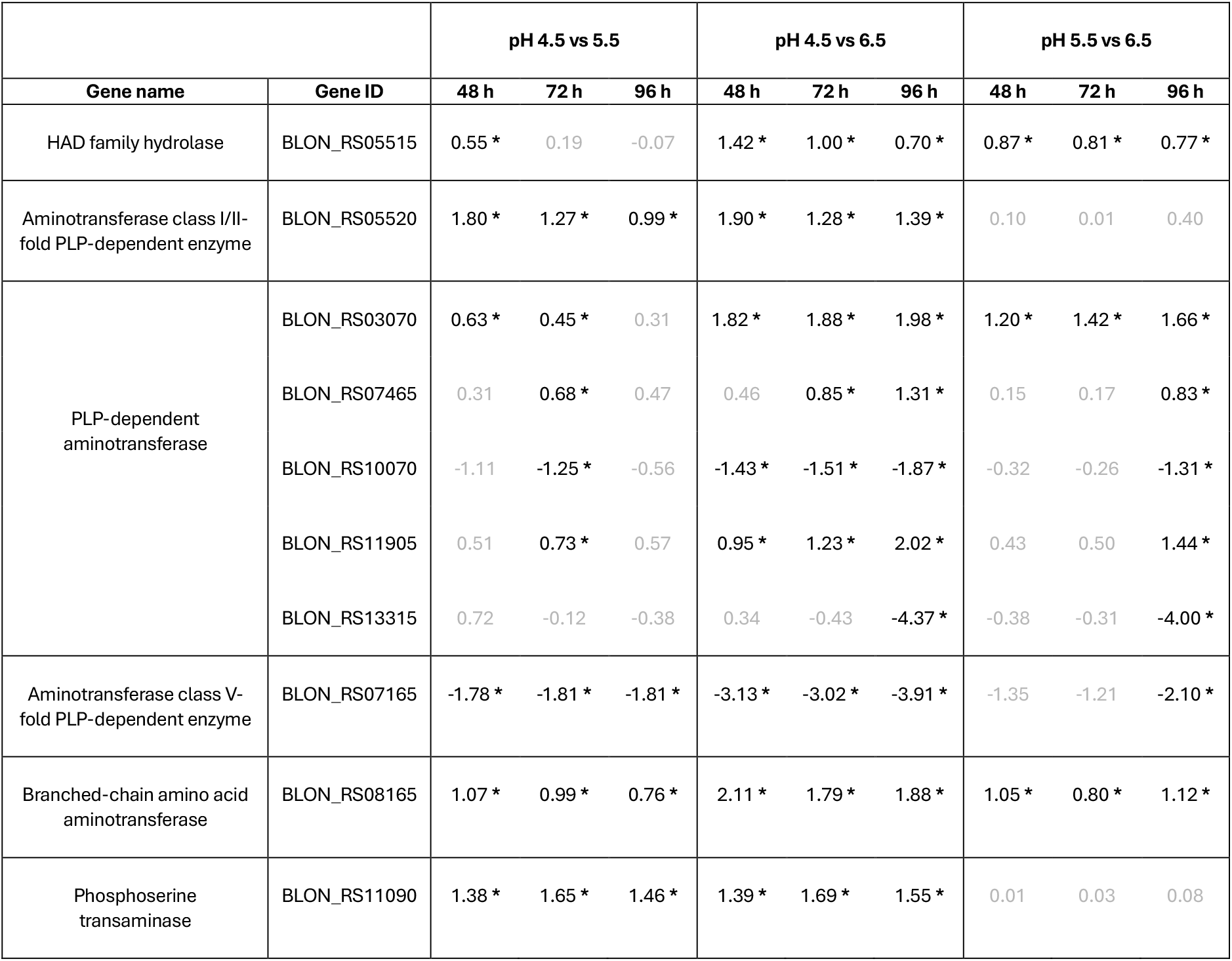
Differential expression of the had gene and genes encoding aminotransferases. The table shows log2 fold changes at each time point, calculated from three biological replicates. *: Genes with an average read count greater than 100 across all samples and an FDR-corrected p-value < 0.1 for the log2 fold change are considered significant.

In *B. breve*, a different *aat* gene has been shown to play a key role in ALA production [23]. The *B. infantis* homolog (BLON_ RS07465) did not show a consistent change in expression when comparing the different pH conditions (**Table 1**). Overall, several genes annotated as aminotransferases were either up- or downregulated in response to low pH (**Table 1**), suggesting that the cells are actively rebalancing amino acid metabolism in response to acid stress.

## Discussion

‘Infant-type’ *Bifidobacterium* species, including *B. infantis*, are key producers of immune-regulating ALAs in the infant gut [5]. However, the factors that determine ALA abundance remain poorly understood. To address this gap, we investigated how environmental factors affect ALA production by *B. infantis in vitro*.

First, we showed that ALA production follows the availability of exogenous AAAs, suggesting that higher dietary AAA levels may lead to increased ALA levels in the infant gut. This observation is consistent with patterns reported in lactic acid bacteria (LAB), where PLA and OH-PLA production increases in response to phenylalanine and tyrosine supplementation [20], [24], and with the dose-dependent relationship between ILA and tryptophan described for *Clostridiales* species [25]. However, it remains to be established if AAA diet supplementation in early life will in fact increase gastrointestinal and circulating ALA levels.

Next, we found that ALA production is tightly coupled to *B. infantis* cell density, indicating that bacterial abundance plays a crucial role in determining metabolite levels. This observation is consistent with reports of growth-associated ALA production in *Lactobacillus* [26], [27] and aligns with findings from the Danish CIG infant cohort, where faecal ALA concentrations positively correlated with the abundance of *B. longum* subsp. *infantis* [5]. Promoting growth of *B. infantis* may therefore be a strategy to enhance ALA production in the infant gut. Reciprocally, it has been suggested that ALA production promotes bacterial growth under nutrient-limited conditions. This has been shown for *B. breve* where supplementing tryptophan and phenylalanine in an otherwise amino acid–limited defined medium had a strong positive effect on growth. Furthermore, mutants in the *aldh* gene did not exhibit the same fitness benefits as wild type *B. breve*, indicating that ALA production is essential for growth under such conditions [23]. However, conditions in the infant gut are unlikely to be amino acid limited, and faecal AAA concentrations are commonly exceeding faecal ALA concentrations [28].

Since high abundance of *B. infantis* is also linked to a low faecal pH [21], we next investigated how pH affects ALA production. We showed that growth is significantly inhibited at pH 4.5 compared to less acidic conditions (pH 5.5 and 6.5), consistent with the weak acid tolerance previously reported for *B. longum* [29]. Consequently, lower absolute ALA levels were observed at pH 4.5. This suggests that a highly acidic gut environment may slow bacterial growth and, in turn, reduce overall ALA levels. Nonetheless, low faecal pH of 4.5-5.0 has been observed in breastfed infants, with gut microbiomes being highly dominated by *B. infantis* [30], indicating that some environmental conditions in the gut reduces its acid sensitivity.

Beyond the growth-mediated effects, pH had a distinct influence on ALA production by altering the ALA profile, favoring PLA production with the expense of ILA at lower pH. This is in line with acid stress triggering transcriptional and proteomic changes in amino acid metabolism in *B. longum* [31], [32]. Since transamination has been described as the rate-limiting step for ALA synthesis in LAB [33], [34], and we observed differential expression of several *aat* genes in response to pH, we speculate that changes in aminotransferase activity may explain this shift in ALA profiles. Supporting this hypothesis, enhancing transamination by increasing the availability of the amino group acceptor α-ketoglutarate (α-KG) has been shown to increase ALA production by *B. infantis*, with PLA exhibiting the most pronounced increase [35]. Furthermore, disruption of the transaminase araT in *B. breve* severely impairs ILA production while having only minor effects on PLA and increasing 4-OH-PLA levels [23]. Since araT is required for the biosynthesis of phenylalanine and tyrosine, this suggests that other transaminases or anabolic pathways compensate by supplying pyruvic acid derivatives for these two amino acids in *B. breve* [23]. To our knowledge, the exact aminotransferase(s) responsible for ALA production by *B. infantis* has not yet been identified, but it can be hypothesized that different routes for aromatic pyruvic acid production exist in *B. infantis* and that these are affected by pH.

Overall, our study shows that ALA production by *B. infantis* is regulated by environmental factors. We propose that ALA production can be enhanced through increased AAA availability and *B. infantis* abundance, and that variations in pH may shape the composition of ALAs produced in the gut of infants.

## Materials and methods

### Bacterial strain and culture conditions

In this study, *Bifidobacterium longum subsp. infantis* DSM20088 (*B. infantis*) obtained from DSM (Deutsche Sammlung von Mikroorganism, Germany) was used. The strain was stored in glycerol stocks at -80 °C until use. Glycerol stocks were streaked on deMan–Rogosa–Sharpe agar (SSI Diagnostica, Cat. No.: 50123) supplemented with 0.05% (w/v) L-cysteine hydrochloride (Sigma, Cat. No.: C7477) (MRS-cys agar plates) and incubated for 48 h at 37 °C in an anaerobic chamber (Don Whitley Scientific Limited) containing 10% H_2_, 10% CO_2_ and 80% N_2_ gases. For overnight cultures, single colonies were resuspended in MRS-cys broth [36], and incubated for 20-24 h at 37°C in the anaerobic chamber.

The MRS-cys broth contained: 10 g/L Peptone from casein (Merck, Cat. No.: 1.07213.1000); 8.0 g/L Meat extract (Oxoid, Cat. No.: LP0029); 4.0 g Yeast extract (Oxoid, Cat. No.: LP0021); 2.0 g/L Di-potassium hydrogen phosphate (Merck, Cat. No.: 1.05104.1000); 5.0 g/L Sodium acetate (Sigma, Cat. No.: S8750), 2.0 g/L Tri-ammonium citrate (Sigma, Cat. No.: A1332); 0.2 g Magnesium sulfate heptahydrate (Merck, Cat. No.: 1.05886.0500); 0.05 g/L Manganous sulfate monohydrate (Sigma, Cat. No.: M7634) supplemented with sterile filtered 0.05% (w/v) L-cysteine hydrochloride (Sigma, Cat. No.: C7477) and 2% (w/v) D-glucose (Merck, Cat. No.: G8270) after autoclavation.

### Batch fermentation with aromatic amino acid supplementation

To test how ALA production by *B. infantis* is affected by AAA availability, *in vitro* fermentations with different initial concentrations of AAA were set up. Singles colonies of *B. infantis* were inoculated into MRS-cys broth supplemented with 0.025, 0.05, 0.1 and 0.2% (w/v) sterile filtered L-tryptophan (Sigma, Cat. No.: T0254), L-phenylalanine (Sigma, Cat. No.: P2126) or L-tyrosine disodium salt dihydrate (Merck, Cat. No.: 1.02413.0100) and incubated in the anaerobic chamber at 37°C. Each AAA supplemented culture was evaluated in biological triplicates and cultures in MRS-cys broth without supplementation served as control. After 48 h of growth, the optical density at 600 nm was measured in a 96-well plate using a PowerWave HT Microplate Spectrophotometer (BioTek). All cultures were centrifuged at 15,000 g for 10 minutes at 4 °C and the supernatants were stored at - 20 °C until LC-MS analysis.

### Chemostat cultivations at different pH levels

To test the influence of pH on ALA production, *B. infantis* was grown in chemostat cultures at a dilution rate of 0.06 h^-1^ in 1 L Biostat A bioreactors (Satorius). The temperature was kept constant at 37 °C, the pH was controlled at constant levels (4.5, 5.5 and 6.5) by automatic addition of 1 M HCl and 2 M NaOH and the stirring speed was 100 rpm. To maintain anaerobic conditions the headspace was flushed with 80% (v/v) N_2_ and 20% (v/v) CO_2_ gases as at rate of 50 ml min^-1^. The chemostat cultures were prepared by adding 500 mL MRS broth to the bioreactors followed by autoclavation of the entire setup. Subsequently, sterile filtered L-cysteine-HCl (0.5 g L^−1^) and D-glucose (20 g L^-1^) were added. After an overnight pre-run to ensure anaerobic conditions, the bioreactors were inoculated 1:100 with an overnight culture made in MRS-cys broth as previously described. After 24 h of growth (batch mode), in- and outflow was applied and continuous fermentation (chemostat mode) was run for further 72 h. Bacterial growth was monitored by measuring the optical density at 600 nm using a spectrophotometer. To determine the concentration of viable cells during the experiment, CFUs mL^-1^ were estimated. Samples were serial diluted in MRS-cys broth and appropriate dilutions were plated on MRS-cys agar plates in duplicates. The plates were incubated at 37 °C in the anaerobic chamber for 48 h, after which colony forming units (CFUs) were counted. For metabolite quantification, samples were centrifuged at 15,000 g for 10 minutes at 4 °C and the supernatants were stored at -20°C until LC-MS analysis. For RNA analysis, one volume of sample was immediately mixed with two volumes of RNAProtect Bacteria Reagent (QIAGEN, Cat. No.: 76506). The stabilized solution was pelleted according to manufacturer’s instructions and stored at − 80 °C until RNA extraction.

### Identification and quantification of aromatic lactic acids by LC-MS

For quantification of aromatic lactic acids and aromatic amino acids, liquid chromatography mass spectrometry (LC-MS) was performed. For the LC-MS, isotopic-labeled L-tryptophan (indole-d5, 98%) (Cambridge Isotope Laboratories) was used as internal standard. To quantify the aromatic compounds, different concentrations of analytes (0 μg ml^−1^, 0.8 μg ml^−1^, 2 μg ml^−1^ and 4 μg ml^−1^) were prepared by mixing analyte mix (**Supplementary Materials Table 1**), internal standard and sterile water. Furthermore, quality control (QC) standards were prepared and processed in the same way as samples to normalize against any loss or gain of the analytes during data processing. The QC standards were prepared in triplicates of 80 μL by mixing growth media 1:1 with analyte mix (each analyte at 40 μg/mL).

To prepare samples for LC-MS, culture supernatants were thawed at 4 °C and then centrifuged at 16,000 g at 4 °C for 10 min. 80 µl supernatant was transferred to a new tube. To each sample and QC standard, 20 µl internal standard (40 µg ml^−1^) and 300 µl acetonitrile (Sigma, Cat. No.: 1003363276) were added. The samples were vortexed for 10 seconds and placed at −20 °C for 10 min to precipitate proteins. Afterwards, the samples were centrifuged at 16,000 g, at 4 °C for 10 min. 50 µl of supernatant of each sample followed by 50 µl of sterile water was transferred to a liquid chromatography vial (resulting in a 1:10 dilution of the sample and an internal standard concentration of 1 µg ml^−1^).

The LC-MS was performed as previously described [16]. In brief, samples (5 µL) were injected and analysed by an UPLC-QTOF-MS system, utilizing Dionex Ultimate 3000 RS liquid chromatograph coupled to a Bruker maXis time-of-flight mass spectrometer with an electrospray interphase (Bruker Daltonics). The analytes were separated on a Poroshell 120 SB-C18 column with a dimension of 2.1x100 mm and 2.7 µm particle size (Agilent, Cat. No.: 685775-902). The LC-MS data from negative electrospray ionization (ESI) mode was processed using QuantAnalysis v.2.2. Peak area ratios were calculated as the integrated area of the metabolite peak divided by the area of the internal standard peak. Standard curves were generated for each metabolite and used to determine its concentration in the samples.

ALAs were measured at the end of fermentation in batch cultures and from 24h and onward in chemostat cultures. In chemostat culture supernatants at pH 4.5, only PLA was measured above 0.8 μg ml−1. However, since extracted chromatograms for both ILA and OH-PLA were identified as clear peaks, extrapolated data were used for quantification.

### Transcriptome analysis

Gene expression in response to pH levels was analysed by RT-qPCR and RNAseq. RNA from chemostat samples was extracted by using RNeasy Mini kit (Qiagen, Cat. No.: 74106) with minor modifications. In brief, 600 µl RLT lysis buffer were added to each sample followed by 10 µl of β-mercaptoethanol to inactivate RNase activity. The samples were then transferred to bead columns and placed in a bead beater (Retch) for 30 seconds at 30 Hz to mechanically disrupt cells. After bead beating, the samples were centrifuged at 8000 g and the supernatant was mixed 1:1 with 80% ethanol. Subsequent extraction steps were performed according to the RNeasy Mini Kit spin column purification protocol. The concentration of extracted RNA was determined using Qubit RNA high sensitivity assay (Invitrogen, Cat. No.: Q32852) and the purity was assessed on the Nanodrop by measuring the A260/A280 and A260/A230 ratios. Samples used for RNAseq, included a DNAse step (Qiagen, Cat. No.: 79254) in the RNA extraction as described in the RNeasy Mini Kit spin column purification protocol.

For RT-qPCR, isolated RNA was reverse transcribed into cDNA using the reverse transcription protocol for SuperScript™ IV VILO™ Master Mix with ezDNase enzyme (Invitrogen, Cat. No.: 11766050).

To normalize *aldh* gene expression, *pdxS* and *gluC* were used as references genes [37]. Primers were designed for *aldh* and reference genes using Primer3web v. 4.1.0 (https://primer3.ut.ee/), considering the following parameters: primer size (18–23 bp), CG% (30-70), primer melting temperature (Tm 55–65 °C) and product size range (100–200 bp). The specificity of the primers was tested *in silico* using BLAST analysis against the genome of *Bifidobacterium longum subsp. infantis ATCC 15697 = JCM 1222 = DSM 20088 (NCBI:txid391904)*. To determine primer efficiencies, qPCR reactions with 10-fold dilutions of genomic DNA (gDNA) extracted from an O/N culture of *B. infantis* using DNeasy UltraClean Microbial kit (Qiagen, Cat. No.: 12224-50) was set up using the method described below. Acceptable primer pairs had PCR efficiencies between 90% and 110%. Primer sequences and information can be found in **Supplementary Materials Table 2**. All qPCRs were performed in 20-µL mixtures containing 10 µL of SYBR Green Master Mix (2X) (Thermo Fisher Scientific, Cat. No.: A25742), 1 µL of each primer (10 pmol/µL), 2 µL of DNA template, and 6 µL of nuclease-free water. qPCRs were performed in 384-Well Clear Reaction plates (Applied Biosystems, Cat. No.: 4483285) (for qPCRs), loaded to a QuantStudio 5 Real-Time PCR system (Applied Biosystems) and were run under the following conditions: pre-incubation at 50°C for 2 minutes, polymerase activation at 95°C for 10 seconds followed by 40 cycles, each at 95°C for 15 seconds, 55°C for 15 seconds and 72°C for 1 minute. Amplification specificity was confirmed by melt curve analysis under the following conditions: 95°C for 15 seconds; 60°C for 1 minute and 95°C for 15 seconds. qPCR reactions were performed in technical triplicates with a non-template control (NTC), RNA extraction control and a no reverse transcriptase control (-RT). 2 ng cDNA was used for qPCR and the relative expression was calculated by the 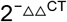 method.

RNAseq was performed by Eurofins genomics (Germany) using Illumina NovaSeq. Raw sequencing data have been deposited in the European Nucleotide Archive (ENA) under accession number PRJEB105461. Raw sequencing data were pre-processed to generate high-quality data for downstream analyses. Reads originating from ribosomal RNA (rRNA) were identified and removed using RiboDetector [38]. The remaining non-rRNA reads were subjected to quality control using fastp [39], including adapter trimming, removal of low-quality bases (Phred quality score <20), and per-read quality pruning using a sliding-window approach. After quality trimming, reads shorter than 30 bp were discarded. For paired-end sequencing, only read pairs in which both reads passed quality control were retained. Sequencing quality was assessed using standard metrics, including the proportion of bases with Phred quality score ≥30 (Q30) and GC content (**Supplementary Table 3**). High-quality reads were aligned to the *B. infantis* DSM 20088 reference genome using STAR (Spliced Transcripts Alignment to a Reference) [40] within the Sentieon framework [41], incorporating known gene models. STAR was used to perform spliced read alignment optimized for RNA-seq data. Gene-level quantification was performed using featureCounts [42]. Low expressed genes were filtered prior to downstream analyses by removing genes with counts per million (CPM) values < 1. The abundance counts of each gene were then used to perform differential gene expression (DGE). DGE was performed using the edgeR package [43] from Bioconductor. Library sizes were normalized using the calcNormFactors() function to correct for differences in RNA composition between samples. Statistical comparisons between experimental conditions were performed for each gene, and p-values were adjusted for multiple testing using the Benjamini–Hochberg false discovery rate (FDR) method. Genes with average read counts greater than 100 across all samples and FDR-adjusted p-values < 0.1 were considered differentially expressed.

### Statistics

All statistical analyses, except for DGE, were carried out in GraphPad Prism software (v. 10.4.1). For the AAA availability test, statistical significance between groups was determined by one-way analysis of variance (ANOVA) followed by Tukey’s multiple comparisons test. For the chemostat experiments, cell densities were compared using repeated measures two-way ANOVA followed by Bonferroni correction for multiple comparisons test. Metabolite differences in the chemostat experiments were compared using two-way ANOVA. Relative percentwise abundance of ALAs were assessed by Chi-square test. Relative *aldh* expression between groups were compared using unpaired two-tailed t-test with Welch’s correction on gene expression values normalized to reference genes at each timepoint. All correlations were Pearson correlations. For all statistical analysis, if not stated otherwise, a p-value < 0.05 was considered as significant.

## Acknowledgements

We thank Anurag Kumar Sinha, Julius Emil Brinck and Emmelie Joe Freudenberg Rasmussen (National Food Institute, Technical University of Denmark) for technical advice concerning chemostat fermentations. Moreover, we thank Bodil Madsen and Katja Ann Kristensen (National Food Institute, Technical University of Denmark) for technical support in the laboratory. This study was funded by a grant from the Novo Nordisk Foundation Challenge program (PRIMA, grant number NNF19OC0056246).

